# Photodegradable Hydrogels for On-Demand Modeling of Age-Related Spatiotemporal ECM Deformation

**DOI:** 10.1101/2025.07.28.667183

**Authors:** Felix Reul, Cédric Bergerbit, Iulia Scarlat, Teodora Piskova, Vasudha Turuvekere Krishnamurthy, Aleksandra Kozyrina, Stacy Lok Sze Yam, Laura De Laporte, Jacopo Di Russo

**Author notes:** Correspondence: Jacopo Di Russo.

## Abstract

The extracellular matrix (ECM) plays a crucial role in regulating tissue behavior through a dynamic interplay of spatial and temporal cues. Dynamic materials capable of modulating these cues at relevant scales are essential for tackling current challenges in tissue engineering and addressing fundamental biological questions. In vision research, there is a notable lack of suitable in vitro systems to study ECM dynamics. To help fill this gap, we developed an easy-to-use, photosensitive poly(ethylene glycol)-based hydrogel that can deform on demand to simulate ECM bulging, known as drusen, associated with the aging of the outer retina. Our findings demonstrate that variations in the size of these artificial drusen during culture impact morphometric parameters of the retinal pigment epithelium, offering new insights into its mechanical resilience to different drusen sizes. Notably, drusen formation in our system does not significantly affect the cellular actin cytoskeleton or polarity, which are often disrupted in conventional acute substrate deformation models, allowing the study of early aging before detectable pathology. In summary, we present a light-tunable hydrogel platform that enables precise spatial mimicry of ECM topographical changes, offering a promising tool for investigating the mechanobiological aspects of dynamic cell-matrix interactions.

## Introduction

The extracellular matrix (ECM) is a fundamental, non-cellular component of all tissues and organs. Beyond providing the structural scaffolding that sustains tissue architecture, it acts as a dynamic regulator of cellular behavior in tissues ^[1–3]^. Tissues continuously evolve through morphogenesis, homeostasis, and aging, with the ECM providing a dynamic array of biochemical, topological, and physical cues ^[1–3]^. Mimicking these dynamic changes at a relevant mechanical and length scale in a laboratory setting is therefore crucial for overcoming current challenges in tissue engineering and addressing basic biological questions. This is evident in advanced stem cell-derived tissues, which closely replicate the metabolic activity, tissue architecture, and developmental timing found in vivo. Unlike conventional, fast-proliferating immortalized cell lines, stem cell-derived systems grow and differentiate in a more controlled and coordinated environment, mirroring the slower pace and complex spatial organization of native tissues^[4,5]^. As a result, recreating the dynamic interplay between cells and the ECM over realistic scales is essential to fully harness the potential of stem cell-derived models for studying fundamental processes, such as tissue remodeling, disease progression, and aging under near-physiological conditions^[4]^.

Vision research faces major challenges due to the lack of relevant in vitro model systems, as animal models often fail to accurately replicate the anatomy, physiology, and disease mechanisms of the human eye. In particular, modeling key age-related changes in the ECM seen in humans remains challenging in animal systems, creating a significant barrier to a comprehensive understanding of the pathophysiology of eye diseases^[6,7]^. One significant age-related change in the ECM of the eye is the local accumulation of lipids and proteins underneath the retinal pigment epithelium (RPE), named drusen. The morphology and composition of drusen differ among individuals. However, epidemiological studies have indicated that medium (63 – 125 µm in diameter) and large (> 125 µm in diameter) drusen, with a height of 17-30 µm, are associated with phenotypical changes and polarity loss in RPE cells. Although drusen of varying sizes are often observed in asymptomatic patients, only their prolonged presence leads to photoreceptor degeneration and contributes to the development of age-related macular degeneration (AMD) ^[8,9]^, highlighting the importance of mimicking and elucidating the underlying cause–effect mechanisms in a model system. Rodent models do not spontaneously develop drusen unless genetically challenged, and even then, they do not fully recapitulate drusen formation and pathogenesis in humans ^[10]^. In contrast, non-human primates remain the gold standard for in vivo drusen research, as they closely recapitulate the molecular features of the human retina. However, in addition to ethical considerations, they have not yet been shown to progress to advanced stages of AMD^[11]^. In vitro modelling provides a valuable alternative, offering a flexible and controllable experimental platform that enables deeper investigation of cause–effect relationships. By allowing precise manipulation of environmental variables, this approach facilitates the distinction between factors that drive healthy versus pathological ageing, an investigation that is far more challenging in an in vivo system. In the past, diverse approaches have been employed to mimic drusen formation and study the consequences on RPE in vitro. Here, techniques such as vacuum, magnetic, piston, or air pressure-induced stretching of the substrate aimed to recapitulate the dynamic appearance of drusen ^[12–14]^. However, these approaches constrain the in vitro system by imposing unphysiological substrate stiffness (MPa range) and fixed topography.

To address this limitation, we revisited the use of photodegradable hydrogels^[15,16]^ as a tissue culture substrate, providing a platform to study the impact of ECM topographical changes on tissues. As a proof of concept, we applied this system to mimic medium- and large-sized drusen and investigated their impact on stem cell-derived RPE. Through qualitative and quantitative analyses, we demonstrated the impact of this methodology, which can induce adaptation of the epithelium on relevant substrate stiffness, suggesting mechanobiological insights with implications for the pathophysiology of retinal degeneration. In conclusion, this work highlights the potential of such a tunable hydrogel system, which can be employed in both two-dimensional and three-dimensional setups to investigate the impact of ECM changes at mechanical and length scales comparable to those in vivo.

## Results and Discussion

### Engineering a phototunable hydrogel to dynamically mimic ECM topographical changes in space and time

As ECM remodeling occurs at specific times and length scales, we aimed to develop a tunable hydrogel platform with a photo-degradable mesh. This would allow us to tune its properties in alignment with the in vivo situations. To achieve this, we took inspiration from the work of Kloxin and colleagues^[15,16]^. We synthesized the photodegradable crosslinker (PDC) containing a linear polyethylene glycol (PEG) core, ortho nitro-benzyl groups as photolabile moieties, and acrylate end groups. The PDC was incorporated in PEG-acrylate-based network through a free radical polymerization initiated by a redox reaction of Ammonium persulfate (APS) and Tetramethylethylenediamine (TEMED) (Figure 1A). The ratio of the different hydrogel precursor components determines the final material properties of the hydrogel, allowing for the tuning of substrate stiffness to mimic in vivo ECM mechanical properties. Furthermore, the ratio of PDC and non-degradable PEG components sets the sensitivity of the hydrogel to light. In fact, ortho nitro-benzyl groups are excited by near-UV light (∼275-405 nm)^[15,16]^, and undergo photolysis, resulting in bond breakage at the benzyl positions, which leads to a partial reduction in the crosslink density within the hydrogel structure. As a consequence, an influx of water will result in swelling of the hydrogel (Figure 1A, B). The simplicity of its implementation is exemplified using a standard inverted microscope, equipped with a 385 nm light-emitting diode (LED), with adjustable intensity and exposure time. Furthermore, the illuminated patterns can be modified by adjusting the stage position, the objective magnification, and the degree of diaphragm opening located between the LED and the objective revolver (Figure 1B). Interestingly, we found that localized illumination causes local swelling, which in turn leads to topographical changes of the hydrogel surface. These changes can be produced in specific patterns, such as arrays (Figure 1C), can be tuned for swelling by varying LED intensity or exposure time (Figure 1D). Furthermore, the illumination-induced swelling is additive regardless of the illuminated area size (100 vs 300 µm in diameter, Figure 1E). Next, curious to understand the effect of such changes on the local mechanics, we illuminated areas with diameters of 100 and 300 µm using an illumination time of 60 seconds with 100 % intensity (3.83 mW cm^-2^). This resulted in surface bulges with an average height of approximately 20 µm (Figure 1F). The stiffness of the resulting bulges was then assessed by nanoindentation using a spherical tip with 20 µm diameter and the Effective Young’s modules was quantified for a cell-relevant indentation depth of 3.5 µm^[17]^. To our surprise, the Effective Young’s modulus for the illuminated areas was comparable to that of both the non-illuminated areas of the photodegradable hydrogel and the illuminated non-degradable control PEG hydrogel (Figure 1G). This is likely because swelling is primarily governed by osmotic fluid uptake and macroscopic network expansion, while the elastic modulus measured at the microscale remains dominated by polymer entanglement and crosslink density close to the surface, which is not fully disrupted during partial degradation^[18]^. Thus, local stiffness can remain unchanged even as bulk deformation increases. In conclusion, we successfully adapted the use of the PDC to create a substrate, allowing for controllable substrate deformations or topographical changes without significantly changing its stiffness.

**Figure 1:**
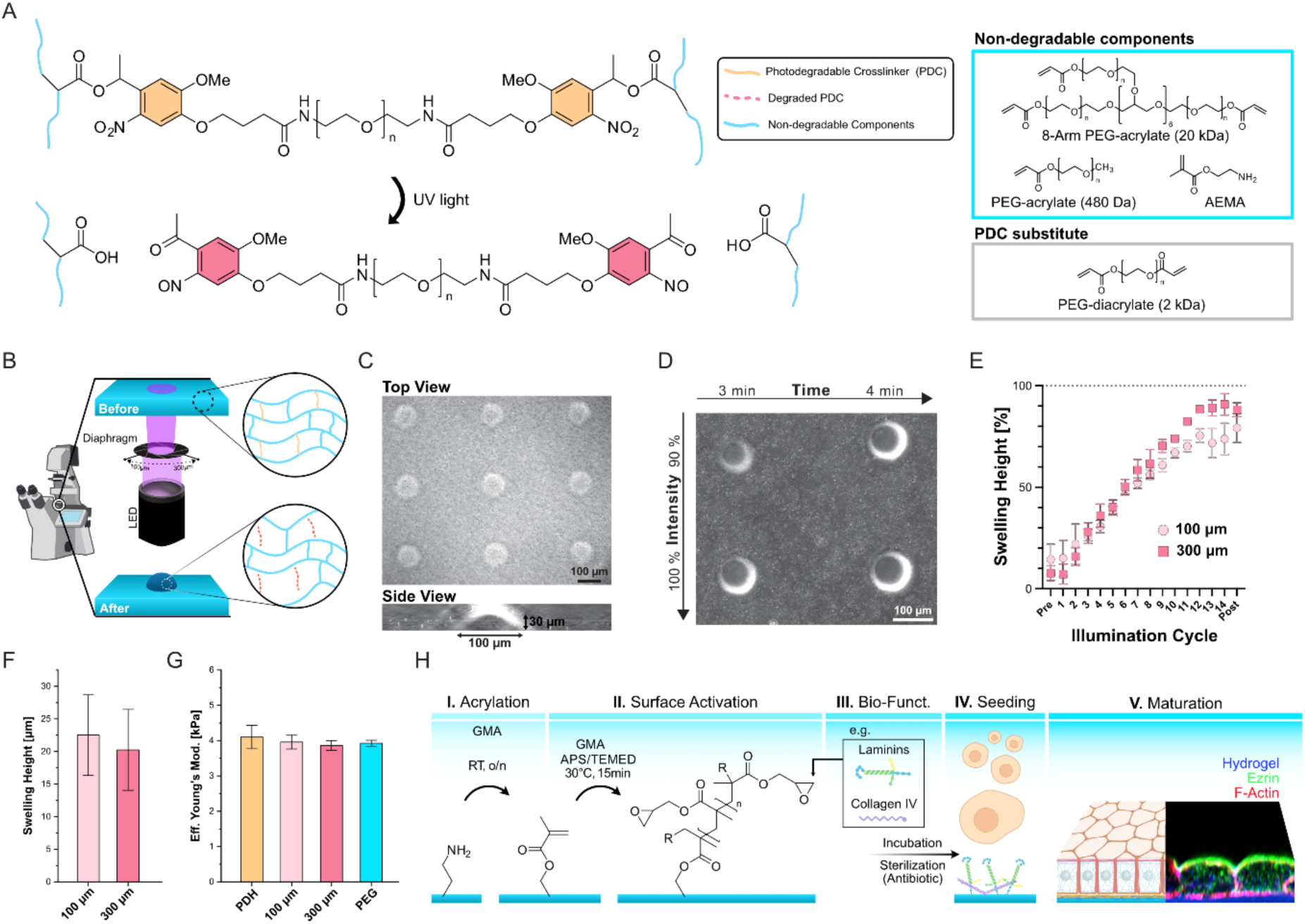
A photodegradable hydrogel to mimic ECM remodeling in relevant time and length scale. **A)** Photodegradable crosslinker (PDC) connected to non-degradable components of the hydrogel undergoes photolysis after illumination with blue (385 nm) light, reducing the crosslinking density. The PDC is incorporated in a non-degradable PEG network composed of 8-Arm PEG-acrylate, PEG acrylate, and AEMA. For a non-degradable hydrogel control, a PEG diacrylate is employed. **B)** Schematic of the experiment setup on an inverted microscope for spatially controlled illumination using an inverted microscope equipped with 385 nm LED and an adjustable diaphragm. **C)** Array patterned to show ease of use and controllability. **D)** Time/Intensity matrix to highlight the potential and versatility of the substrate. **E)** Additive illumination plotted as percentage swelling height ± standard error mean (SEM) over illumination cycles, with each cycle being 4 seconds of UV illumination at 3.83 mW cm^-2^ followed by a break of one hour. The post-experiment reference image was taken 9 hours after the last illumination cycle. Independent repetitions (N): 100 and 300 µm PDH, N=3. **F)** Average swelling height ± SEM of localized illumination of photodegradable hydrogels with a diameter of 100 and 300 µm. Independent repetitions (N): 100 and 300 µm PDH, N=3. **G)** Average Effective Young’s modulus ± SEM of the non-illuminated photodegradable hydrogel (PDH), illuminated with a diameter of 100 or 300 µm, and non-degradable PEG control (PEG). Independent repetitions (N): 100 and 300 µm PDH, N=3. Statistical significance in E and F was assessed using a t-test and one-way ANOVA, respectively. **H)** Illustrated bio-functionalization steps starting after the production of the hydrogels. I) incubation with GMA overnight at room temperature; II) Free radical polymerization of a GMA solution for 15 minutes at 30°C; III) Surface functionalization with a protein mixture overnight at 37°C, followed by sterilization with antibiotic solution; IV) Cell seeding; V) Maturation. The schematic shows an example of a polarized epithelium.

To support cell adhesion and bioactivity, hydrogel surfaces typically require functionalization with ECM proteins or adhesion ligands. In comparison to common cell culture substrates such as plastic surfaces, glass, or polydimethylsiloxane (PDMS), where protein can mostly be deposited by physisorption, the bio-functionalization of PEG hydrogels is primarily achieved by chemisorption using UV-initiated surface activation^[19–21]^. Since photodegradable hydrogels cannot utilize UV-driven post-functionalization strategies, we developed an alternative strategy. This strategy focuses on an orthogonal reaction route to address the lack of free acrylates after gel polymerization, which are necessary for secondary functionalization reactions. 2-aminoethylmethacrylamide (AEMA) was copolymerized into the polymer backbone, resulting in free amine groups that can be exploited in a post-functionalization step (Figure 1H). The post-functionalization was achieved by the addition of glycidyl methacrylate (GMA) to access the free amines in the hydrogel and convert them into meth-acrylate groups through an amine-epoxy ring-opening reaction (Figure 1H, step I). These new acrylates can be further accessed through free radical polymerization with GMA and the branching agent tetramethyl acrylate to build short amine-reactive polymer chains on the surface of the hydrogel (Figure 1H, step II). Finally, the hydrogel can be incubated with a mixture of the desired adhesive proteins (e.g., basement membrane proteins such as laminins and collagen IV) for 24 hours at 37°C in a standard cell culture incubator. After a sterilization step with an antibiotic mixture, the gels can be successfully used to support cell adhesion and maturation (Figure 1H, steps III-V).

Overall, we have now optimized a production and coating strategy of a light-tunable platform to mimic and study the impact of ECM deformation on a given biological system in vitro.

### The photodegradable hydrogel enables the controlled investigation of age-related ECM deformation in a stem cell-based RPE model

To demonstrate the applicability of our photodegradable hydrogel system, we aimed to investigate the impact of substrate topographical deformation on stem cell-derived RPE as a model for studying the correlation between drusen size and the RPE phenotypical switch. We recently demonstrated that this model reliably investigates RPE mechanobiology, as it recapitulates key features of mature human tissue, including its polarization and postmitotic nature^[22,23]^. Stem cell-derived RPE were able to form continuous monolayers on flat photodegradable hydrogels of 4 kPa in stiffness, coated with a mixture of basement membrane laminins and collagen type IV. This setup recapitulates both the biochemical (coating) and mechanical (stiffness) properties of adult Bruch’s membrane, as characterized in a prior work^[22]^. The photodegradable hydrogels were then illuminated to induce local substrate bulging of a diameter of 100 or 300 µm to simulate medium- and large-sized drusen^[8]^ (Figure 2A). To generate the artificial drusen, selected areas were illuminated for 60 seconds at 3.83 mW cm^-2^. The illumination caused a nearly immediate swelling, which reached a plateau only after 8 hours, independent of the size of the illuminated area (Figure 2B). To assess the potential impact of UV phototoxicity, RPE monolayers were obtained on PEG hydrogels synthesized using the PDC substitute and also exposed to UV light for 60 seconds at 3.83 mW cm^-2^. To assess apicobasal polarity of RPE, we stained for Ezrin, a specific marker for apical microvilli^[24,25]^. After 24 hours following drusen induction, qualitative evaluation of high-resolution staining revealed no significant differences between cells on artificial drusen and those in control samples (Figure 2C). This result indicates that neither deformation nor UV illumination significantly impacted cell polarity, and any observed variations were attributable to normal biological heterogeneity. Next, we looked at actin cytoskeleton organization revealing that cells directly atop the artificial drusen showed no obvious morphological differences compared to control regions, with normal actin belt and cortical distribution. However, a close examination of the orthogonal projections of the monolayers suggested changes in monolayer height across the artificial drusen region, both 100 µm and 300 µm in size (Figure 2D).

**Figure 2:**
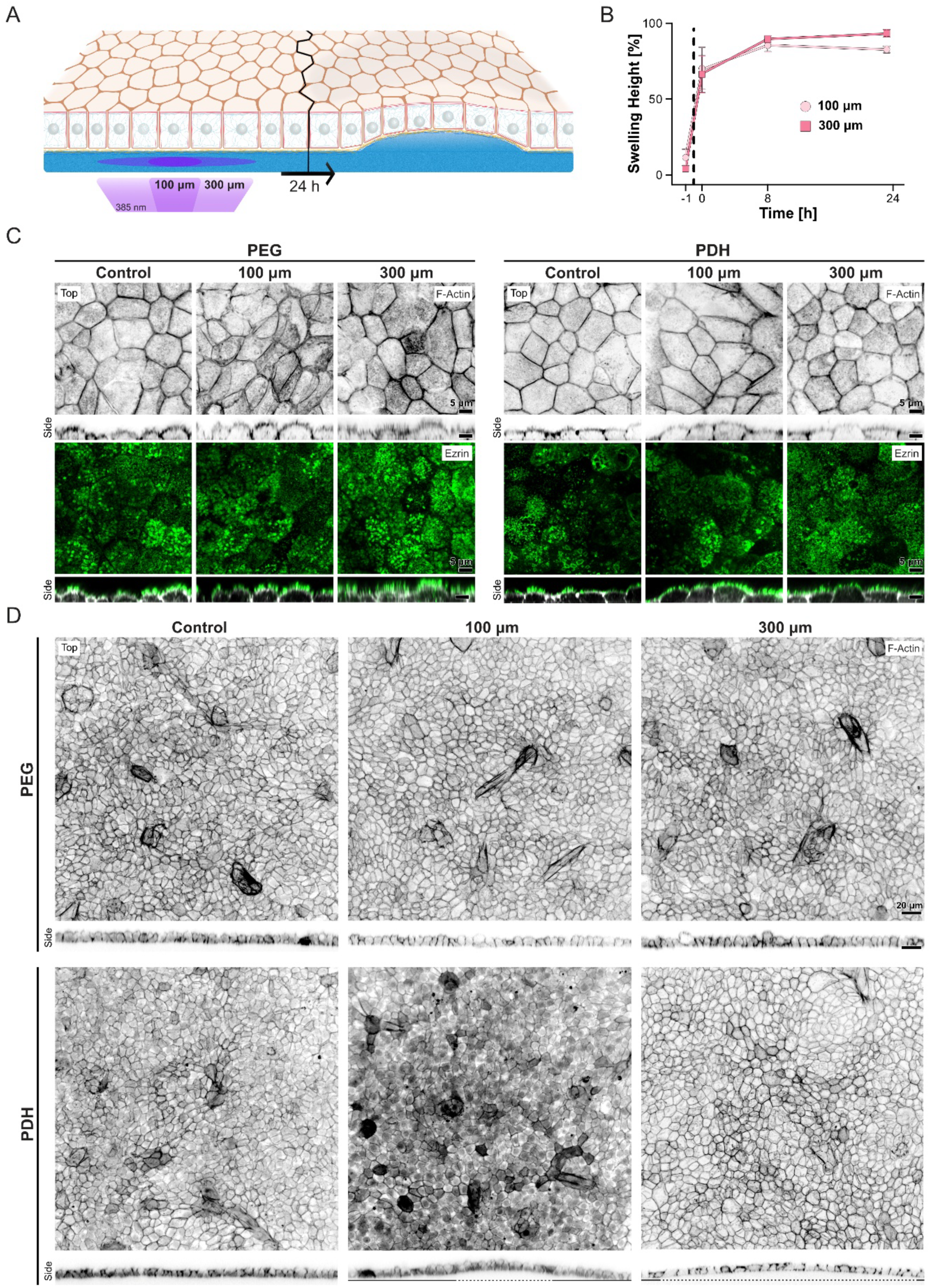
Qualitative characterization of RPE adaptation to artificial drusen at monolayer scale. **A)** Schematic overview of the dynamic formation of an artificial drusen. **B)** Percentage swelling height ± SEM of the artificial drusen over time. The dashed line between t=-1h and t=0h indicates the induction of artificial drusen by UV illumination. Independent repetitions (N): 100 and 300 µm PDH, N=3. **C)** Representative images of top and orthogonal side views of RPE on illuminated and non-illuminated controls and artificial drusen stained for F-actin and Ezrin. **D)** Overviews of RPE monolayers stained for F-actin and orthogonal side views obtained on photodegradable hydrogels (PDH) or PEG hydrogels, with or without illumination of a 100 µm and 300 µm diameter area. Dotted lines indicate the diameter of the illuminated regions.

To quantitatively compare tissue-scale implications of artificial drusen induction, the obtained bulges in the monolayers were analyzed in three distinct regions (Figure 3A). The first region, termed "*Top*" (T), extended radially from the center up to 25 μm for both conditions. The second region, called "*Edge*" (E), was defined as extending 25 µm radially inwards from the respective illumination diameters on the artificial drusen. Lastly, the “*Outside*” (O) region was selected between 150 and 175 µm from the center of the artificial drusen, ensuring that it was not influenced by the surface deformation. Orthogonal views of F-actin-stained RPE monolayers were analyzed to determine monolayer thickness across different regions. The apical and basal cell sides were identified by the intensity of signal in line plots (Figure 3B). The normalized results showed that 100 µm drusen resulted in an average increase in RPE height of approximately 5% at the *Top* and 3% at the *Edge* regions compared to the *Outside* areas, non-illuminated control, and illuminated PEG control. In contrast, the presence of 300 µm large artificial drusen led to a decrease in RPE thickness of about 5% at the *Top*, while the *Edge* region remained unchanged compared to the *Outside* and controls (Figure 3C). Overall, this means that while the cells on top of the 100 µm artificial drusen are taller, in the same region on 300 µm artificial drusen, the cells become shorter, suggesting that medium-size drusen “relaxes” the RPE monolayer, whereas the larger drusen “stretches” the monolayer at the Top region.

**Figure 3:**
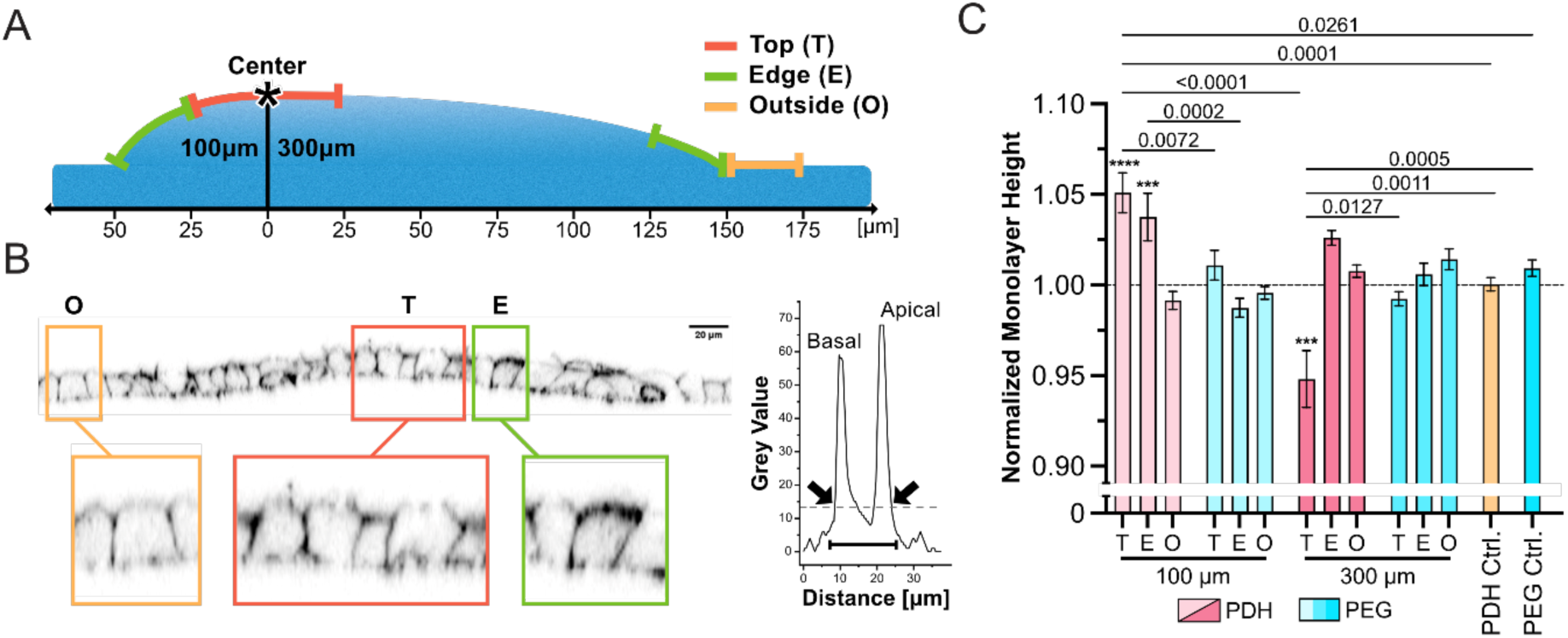
Quantitative characterization of RPE monolayer thickness adaptation to artificial drusen. **A)** Schematic representation of the division of the regions from the center of the illuminated region (*) employed in the analyses. 100 µm illumination is divided into *Top* 0-25 µm and *Edge* 25-50 µm. 300 µm illumination is divided into *Top* 0-25 µm and *Edge* 125-150 µm. For both conditions, the *Outside* region is defined as 150-175 µm. **B)** Representative orthogonal view of a 100 µm artificial drusen with three regions highlighted. The profile of the line plot shows the grey value over the distance. Two peaks indicate the basal and apical actin signal. The dashed line indicates the average grey value of the profile, and the arrows show the points the script recognizes as apical and basal delimitators. Finally, the monolayer height is determined as the difference between the two points. **C)** Bar plot showing the average RPE height ± SEM on four illuminated hydrogels in the three regions *Top (T)*, *Edge (E)*, *Outside (O)* and the two controls *Outside* regions. Data were normalized to the average monolayer height of each respective sample. Statistical analysis was performed using unpaired two-way ANOVA with Tukey’s correction for multiple comparisons. p-values corresponding to the comparison are indicated on the graph. *** indicate a of p-Value= 0.0008 and **** a p-Value = <0.0001 when compared to the relative *Outside* region. Independent repetition (N): 100 µm PDH and PDH control, N=5; 100 µm PEG, N=4; 300 µm PDH, 300 µm PEG and PEG Control, N=3.

Next, we quantified the average cell area and shape factor (P/√A) in the different regions. The shape factor is a metric used to characterize the mechanical state of epithelial tissues, where higher values reflect an increased influence of cell-cell adhesion and unjamming, whereas lower values reflect greater cortical tension and a jammed homeostatic state ^[26]^. The lateral outline of the cells was determined by the maximum projection of the tight junction signal ZO-1 (Figure 4A, Supplementary Figure 1A). A comparison of the average cell areas and shape factors revealed no significant differences between the conditions (Supplementary Figure 2), indicating that the monolayers on the drusen were still jammed in a homeostatic state. Nevertheless, the data showed notable variability and emerging trends in cell area, suggesting the presence of different subpopulations. To investigate this further, we analyzed the frequency distribution of the quantified values. Interestingly, the frequency curve of cell areas for the 100 µm artificial drusen revealed a bimodal behavior for the cells at the *Top* region, a pattern not observed for the *Top* region of the RPE on 300 µm large artificial drusen (Figure 4B and C). In fact, a more unimodal distribution was found in this condition, in the controls, and at the *Edge* and *Outside* areas of both drusen sizes (Supplementary Figure 3). The difference in area distribution supports the hypothesis that a change in the organization of the monolayer may be caused by the disruption of an existing force balance, which induces a topological shift in RPE on 100 µm artificial drusen, but not on 300 µm artificial drusen.

**Figure 4:**
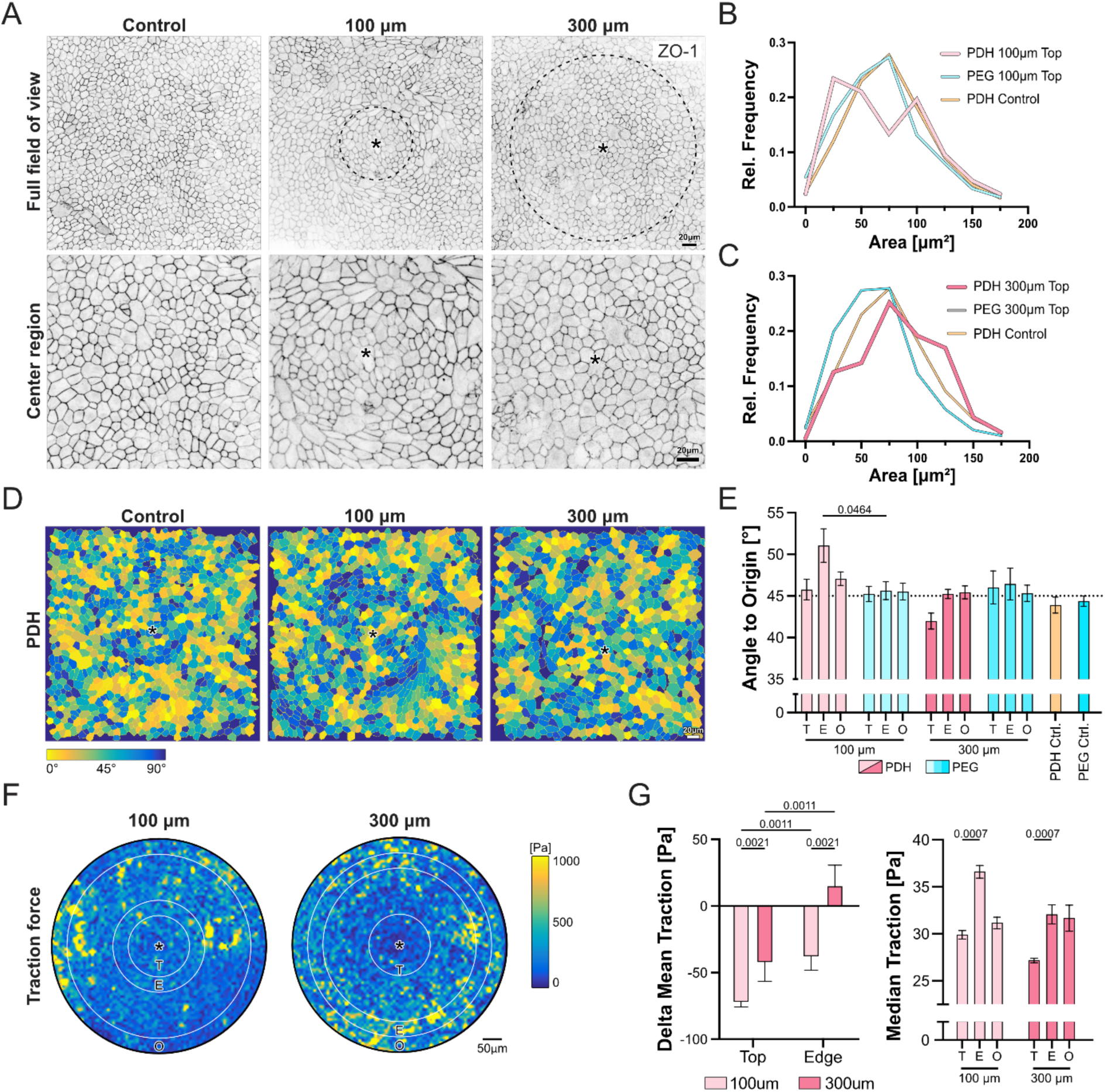
Morphometric characterization of RPE adaptation to artificial drusen at a cellular scale. **A)** Representative overviews (top) and magnifications (bottom) of RPE monolayers stained for ZO-1 on photodegradable hydrogels (PDH) with or without illumination. The illuminated areas are indicated by a dashed line. **B, C)** Relative frequency distributions of the cell areas for *Top* regions of 100 µm (B) and 300 µm (C) artificial drusen, non-illuminated control. **D)** Representative color-coded maps of angular orientation of RPE cells with respect to the drusen center. In yellow is indicated 0° angle towards the center of artificial drusen (or center of the picture for the control), and in dark blue 90° angle, indicating a perpendicular orientation (Left). **E)** Bar plot of the average cell orientation angle to origin ± SEM quantification for the four illuminated conditions and non-illuminated controls (Right). Statistical analysis was performed using unpaired two-way ANOVA with Tukey’s correction for multiple comparisons. Independent repetition (N): 100 µm PDH, 300 µm PDH and PEG Control, N=3; 100 µm PEG, N=6; 300 µm PEG, N=5; PDH Control, N= 4. **F**) Representative traction force plot of 100 µm and 300 µm artificial drusen with the center of the respective artificial drusen marked by an asterix and the *Top*, *Edge* and *Outside* regions outlined. **G**) Bar plot of the delta of the mean traction ± SEM of 100 and 300 µm artificial drusen in relation to the *Outside* region (Left). Bar plot of the median traction ± SEM of 100 and 300 µm artificial drusen (Right). Statistical analyses were performed using unpaired two-way ANOVA with Tukey’s correction for multiple comparisons. Independent repetition (N): 100 and 300 µm PDH, N=3 (Right).

Following this logic, we analyzed cellular orientation as a proxy for assessing the mechanical continuum in epithelia, given that epithelial cells typically align with maximum average stress ^[27–29]^. We quantified the relative orientation of the cells in relation to the center of the artificial drusen by calculating the scalar product between the absolute angle of the cell and the angle directed towards the center of the artificial drusen (Supplementary Figure 1B). This angle varies from 0° to 90°, indicating alignment towards the drusen’s center or orthogonal positioning, respectively (Figure 4D). While we could not observe any clear orientation pattern in RPE cells on 300 µm drusen and controls, we identified a significant circular orientation along the 100 µm drusen *Edge* region (Figure 4D and E). This response can be described as a ring forming around the artificial drusen, separating the inner *Top* region from the *Outside* region. In contrast, for the 300 µm artificial drusen, we observed a less prominent response in the *Top* region, where cells showed a tendency to align towards the center.

Motivated by the observed differences in monolayer height and cell orientation, we performed traction force microscopy to quantify the adhesion strength of RPE cells to the substrate. Quantification and comparison with the background traction in *Outside* regions revealed a reduction in traction forces over both the *Top* and *Edge* areas of RPE monolayers on 100 µm drusen. In contrast, on 300 µm drusen, the reduction in traction was less pronounced on the top region, while the edge regions exhibited a slight increase (Figure 4F and G). Analysis of median traction forces further highlighted distinct regional patterns between 100 µm and 300 µm drusen. On 100 µm drusen, RPE cells displayed significantly higher traction at the edges, consistent with the observed orientation differences, whereas on 300 µm drusen, traction medians were reduced only on the *Top* and remained stable at the *Edges* (Figure 4G). Together, traction force analyses indicate that, regardless of drusen size, RPE cells on drusen exhibit overall reduced traction. However, only RPE monolayers on 100 µm drusen show a distinct *Edge* response, reflected by the shift in median traction, suggesting a discontinuity in the monolayer between the *Top* and surrounding regions.

Collectively, our data showcase the importance of spatial control in mimicking ECM topographical deformation when studying RPE phenotypical changes. In contrast to more acute in vitro systems^[12,14]^, here the RPE monolayers adapt to ECM bulging in a manner more akin to in vivo conditions, with the early appearance of drusen, without the immediate and extreme loss of polarity or integrity seen with acute or non-physiological deformations. Instead, the quantified differences in mechanical stress on 300 µm artificial drusen, which are inferred by monolayer thinning, may provide mechanobiological insights into the pathophysiology of the RPE phenotypical switch observed in vivo.

## Conclusion and Outlook

In this study, we optimized a photodegradable hydrogel that can be tailored to achieve physiologically relevant stiffness and on-demand dynamic deformation. Additionally, we developed a bio-functionalization protocol that permits cell adhesion without relying on UV light. Our results demonstrated successful cell attachment and the formation of a monolayer of stem cell-derived RPE cells. Furthermore, we were able to induce dynamic changes in the topography of the substrate on demand, with variations in two different sizes, simulating ECM deformation (drusen) as a characteristic of retinal aging. By using a mechanically relevant substrate and inducing osmosis-driven deformations, our system enables investigation of a progressive, age-mimicking response distinct from acute reactions.

Using this system, we found that the multicellular response varies with the size of the induced swelling, resulting in distinct differences in monolayer adaptation in terms of height and cell morphology, specifically cell area and orientation, and traction forces without altering cellular polarization, as highlighted by Ezrin staining. These findings suggest that larger deformations are creating higher mechanical stress on the monolayer, which can eventually lead to the loss of cellular polarity in combination with other environmental stresses. On the other hand, medium-sized deformations induce cellular reorganization that may protect cells from mechanical stress and loss of polarity. Interestingly, this is consistent with the higher probability of large drusen to induce retinal degeneration compared to medium-sized drusen ^[8,9]^. Specifically, smaller disruptions are less likely to induce retinal degeneration; however, crossing an unknown threshold appears to have more significant effects on retinal health and vision.

This work has yielded exciting new insights into a challenging-to-study pathology and can serve as a starting point for further research on mechanobiology aspects of retinal degeneration. Most importantly, the deformations of the hydrogel, which preserve the relevant substrate stiffness, do not dramatically disrupt monolayer polarity, as has been observed in other substrate deformation models ^[12–14]^. Although existing in vivo and in vitro models study the connection between drusen and RPE phenotypic changes, they have yet to systematically consider the influence of drusen size, environmental mechanical factors, or the differences between healthy and pathological age-related RPE changes. Notably, in our proof-of-concept study, we did not observe significant changes in polarity in the RPE monolayer. Instead, we detected structural and organizational alterations with clear mechanobiological consequences. These findings highlight the potential of our hydrogel model for studies that combine ECM deformation with other age-related stressors, such as oxidative stress or genetic predisposition, to more comprehensively investigate the concomitant factors that drive the transition from healthy to pathological ageing. Moreover, the versatility of the photodegradable hydrogel enables future research to address the consequences of drusen formation over more extended periods (for example, through multiple illumination events) and changes in their density. This approach may provide deeper insights into the adaptation strategies of RPE observed in vivo ^[30]^.

In conclusion, this photodegradable hydrogel has potential for diverse model systems, as its initial bulk stiffness and protein functionalization can be precisely tuned to replicate the ECM of the target tissue. Moreover, its post-illumination mechanical properties and topographical features can be finely adjusted by modifying the hydrogel composition. The ability to control illumination patterns adds another level of versatility, enabling researchers to dynamically mimic ECM remodeling in both time and space at length scales relevant to tissue developmental, age-related, and mechanobiological processes.

## Methods

### Photodegradable crosslinker synthesis

The synthesis of the photodegradable crosslinker (PDC) followed the published work from Kloxin et al.^[16]^. Briefly, under an inert atmosphere, 4-(4-(1-Hydroxyethyl)-2-methoxy-5-nitrophenoxy)butanoic acid (Iris Biotech) was dissolved in dry dichloromethane (Carl Roth, No.1A9K.1) and deprotonated by triethylamine (Sigma Aldrich, 471283). After cooling the reaction mixture in an ice bath, Acryloyl chloride (Sigma Aldrich, A24109-100G) was added dropwise. After complete addition, the ice bath was removed, and the reaction mixture was stirred at room temperature overnight, then poured into water and a liquid-liquid extraction with chloroform was performed. The organic phase was dried to yield 4-(4-(1-(acryloyloxy)ethyl)-2-methoxy-5-nitrophenoxy)butanoic acid. Its purity was confirmed using nuclear magnetic resonance analysis. In the final step of the reaction, 1-Hydroxybenzotriazol hydrate (HOBt - Sigma Aldrich, 54802-10G-F) and 2-(1H-Benzotriazole-1-yl)-1,1,3,3-tetramethylaminium hexafluorophosphate (HBTU - Sigma Aldrich, 8510060025) were dissolved in N-Methyl-2-pyrrolidone (Sigma Aldrich, 443778), then activated with N, N-Diisopropylethylamine (Sigma Aldrich, 387649). After 5 minutes, PEG-bis amine (Biopharma PEG, HO005005-2K) in N-Methyl-2-pyrrolidone was added, and the mixture was stirred for 24 hours. The reaction mixture was then precipitated in cold diethyl ether, centrifuged, and the resulting brown paste was resuspended in water and dialyzed against five liters of water for three days, with the water changed twice daily. After lyophilization, the product was obtained as a brown solid and purity was checked by nuclear magnetic resonance and electrospray ionization mass spectrometry.

### Hydrogel synthesis

To covalently bond the gel to the glass surface the glass bottom dishes (Cellvis, D35-20-0-N) were activated with 3-(Trimethoxysilyl)propylmethacrylat (abcr, AB109006) with an adapted protocol from Sigma Aldrich (Sigma Aldrich, M6514). Here repeated in short: The glass surface was activated by O_2_ plasma for 1 min at 200 W in a plasma oven. A 3- (Trimethoxysilyl)propylmethacrylat ethanol solution was mixed with diluted acetic acid to allow methanol as a leaving group after the nucleophilic attack of the activated glass surface to the silane. The activated reaction mixture was transferred into the glass bottom dish and the surface functionalization proceeded for three minutes. Afterwards, the dishes were washed with distilled water and ethanol and put in the oven for 15 min at 60°C to evaporate the remaining washing solvents.

PDMS rings with an inner diameter of 4 mm and a height of 200 µm (Darwin Microfluidics, DM-S-PDMS-200) were placed on activated glass bottom dishes. The gel solution was prepared by sequential addition of the components. For the photodegradable gels with an Effective Young’s modulus of 4 kPa the composition was as follows: Poly(ethylenglycol)methyletheracrylat (10 µL, 100 mM, 556 eq., Sigma Aldrich, 454990-250ML), 2-Aminoethylmethacrylat -hydrochlorid (10 µL, 7.55 mM, 5 eq., Sigma Aldrich, 516155-5G), PDC (22.5 µL, 75 mM, 112 eq.), 8-Arm PEG-Acrylate (1.5 µL, 10 mM, 1 eq., Biopharma PEG, A88009-20K), Ammonium persulfate (APS) (4 µL, 1.97 M, Sigma Aldrich, 248614-5G) and lastly N,N,N′,N′-Tetramethylethylenediamine (TEMED) (2 µL, 1.98 M, Sigma Aldrich, 411019-100ML). All the monomers amounted to 17.6 wt/vol%. The composition for the non-degradable gels with an effective Young’s modulus of 4 kPa the composition was adjusted to: Deionized water (20 µL), Poly(ethylenglycol)methyletheracrylat (10 µL, 100 mM, 556 eq.), 2-Aminoethylmethacrylat -hydrochlorid (10 µL, 7.55 mM, 5 eq.), Poly(ethylene glycol) diacrylate (22.5 µL, 100 mM, 150 eq. Sigma Aldrich, 701971-1G), 8-Arm PEG-Acrylate (1.5 µL, 10 mM, 1 eq., Biopharma PEG, A88009-20K), APS (1 µL, 1.97 M), TEMED (0.5 µL, 1.98 M). After the addition of APS and again after the addition of TEMED the reaction mixture was vortexed and shortly spun down to ensure homogeneous distribution of all the monomers and initiators. Roughly 15 µL gel solution was pipetted into the PDMS rings, covered with a Teflon disc, and transferred onto a heater set to 30°C for temperature-controlled polymerization for 1 hour. Afterwards, the gels were flipped upside down for an additional 20 min to ease Teflon-disc detachment, which was promoted by 5 min sonication. The hydrogels were immediately used for the next steps or stored in the dark for a maximum of 12 hours in distilled water at room temperature.

### Nanoindentation

The Effective Young’s modulus of the photodegradable and non-degradable gels was measured by nanoindentation, using a Chiaro Nanoindenter (Optics 11 Life). The probe had a 20 μm spherical glass tip and 0.025 N/m cantilever stiffness. The samples were indented using displacement mode with a maximum depth of 3.5 μm. Analysis of the indentation data was performed in the Optics 11 analysis software (DataViever V2, version 2.6.3) following the Hertz contact model fitting while using single fit mode up to 100% of Pmax and filtering values for a R^2^ greater than 0.95.

### Hydrogel functionalization

The hydrogels were incubated with a Glycidyl methacrylate (GMA) solution (75 μM, Sigma Aldrich 151238-100G) overnight. After removing the unreacted GMA the functionalization solution consisting of: 590 µL H2O, 3uL GMA, 10 µL Di(trimethylolpropane) tetraacrylate (TMA) (0.2 vol/vol% in EtOH, Sigma Aldrich, 408360-100ML), 10uL APS (1.97 M), 2.5 µL TEMED (1.98 M) was added on top of the hydrogels and the gels were put on a heating plate at 30°C, 15min for photodegradable gels and 10 min for non-degradable gels. Directly afterwards the excess poly-GMA was washed off and one by one a hydrophobic barrier with an inner diameter of 6 mm was placed around the gel and immediately the protein solution consisting of: 38.5 µL PBS, 7.5 µL Laminin-332 (100ug/mL, Biolamina, LN332), 7.5 µL Laminin-511 (100 ug/mL, Biolamina, LN511) and 1.5 µL Human Collagen type IV (30 µg/mL, Sigma-Aldrich, C7521-10MG) was pipetted onto the gel. Lastly, the dishes were placed in a cell culture incubator overnight and before use sterilized by washing twice with gentamicin solution for 15 min. Hydrophobic barriers with an inner diameter of 4mm were placed around the sterilized gels and a bubble of 30 µL cell culture medium was placed on top of the gels.

### Cell culture

Before seeding the human induced pluripotent stem cell-derived RPE cells (Fujifilm Cellular Dynamics, R1102) on the hydrogels, the culture was expanded for a week on vitronectin XF (2.5 µg/mL, 440µL, StemCell technologies, 07180) coated 24-well plates as previously reported^[22]^. The medium composition followed a scaled up version of the manufacturer’s instructions; in short it contained MEM-alpha with nucleosides (273.9 mL, Thermo Fisher, 12571063), 100x Glutamax (2.79 mL, Thermo Fisher, 35050061), KnockOut Serum replacement (15 mL, Gibco, 10828028), N2-Supplement (3 mL, Gibco, 17502048), Hydrocortisone (0.33 mL, 50 µM, Sigma Aldrich, H6009), Taurine (1.5 mL, 50 µg/mL, Sigma Aldrich, T8691), Triiodo-L-thyronine (0.21 mL, 20 ng/mL, Sigma Aldrich, T5516) and Gentamicin (0.15 mL, 50 mg/mL, Gibco, 15750060); and medium changes were performed every two days. To generate a single cell solution necessary for seeding on hydrogels the cells were detached by 30 min incubation with 5 mM PBS/EDTA (Sigma Aldrich, E6511), followed by a gentle mechanical wash and finished by a 5 min incubation with TrypLE Express (Gibco, 126050). A final concentration of 2.5 million cells/mL was set and 30µL of cell suspension was used per gel. The cells attached overnight and on the next day 1.5 mL of medium was added to the dish. The culture was maintained for a week with two medium changes.

### Artificial drusen formation

The illumination was performed on a Zeiss Axio Observer 7 with incubation and CO_2_ buffering. Using a 20x objective and a LED diode with 3.83mW cm^-2^ at 100% intensity of the 385 nm channel. The size of the artificial drusen was regulated by opening or closing the diaphragm of the Zeiss Axio Observer and calibrated for each condition. Multiple positions were set, ensuring sufficient spacing to prevent overlap of interference between regions. Additionally, the focal plane was set to 50 µm below the cell monolayer, within the gel. Afterwards, the positions were illuminated one by one for 60 s (230 mJ cm^-2^) and the gels were put back into the incubator. Fixation was performed 24 hours after illumination with 2 w/v% PFA in PBS (Thermo Fisher, J61899.AK) at room temperature for 8 min. Swelling rates were measured by acquiring a reference Z-stack with the focal plane centered on the hydrogel surface. Immediately after UV illumination at all artificial drusen positions, a second Z-stack was acquired, and the change in height was determined from the brightfield focus plane at the top of each forming structure. This measurement was repeated at 8 and 24 hours post-illumination, and the height values for each artificial druse were normalized with 0 indicating the minimum and 1 the maximum.

### Quantification of additive illumination

PDH substrates containing fluorescent beads (0.1 µL of 50% solution, Invitrogen FluoSpheres™ 0.1 µm orange, 540/560) were imaged and illuminated using the Axio Observer 7 microscope. The red fluorescence channel was used to capture images throughout the complete Z-stack (40 slices, 4 µm step size, with the focus plane on the gel surface), while the blue channel was restricted to illuminate only the 8th slice, corresponding to 12 slices (48 µm) from the gel surface. A 1-hour break was included between cycles to allow the hydrogel to equilibrate. The imaging cycle was repeated 15 times, resulting in a total additive illumination time of 60 seconds. A reference image was acquired 24 hours after the first cycle.

### Immunofluorescent staining and imaging

The fixed samples were prepared for immunofluorescent staining by incubation with Triton-X-100 (0.3 vol/vol%, Sigma Aldrich, T8787) for 2 min at room temperature. After washing 3 times with PBS unspecific binding of the antibodies was prevented by 30 min 1% BSA (Serva, 11930) blocking step. With this, sample preparation was finished and the primary antibodies in blocking solution were added for overnight incubation at 4°C. To remove unbound primary antibodies, the samples were washed three times with PBS before addition of secondary antibodies and incubation for a minimum of 2 hours at room temperature. Lastly, the samples were washed again and finally stored in PBS prior to imaging. Samples were imaged at a Zeiss LSM710 Confocal Microscope with a 40x water immersion objective (Zeiss, 420862-9970-000) in normal confocal mode (Z-stack 1.1 µm optical slicing) and Airyscan super resolution mode (Z-stack 0.25 µm optical slicing). Image processing and quantification were performed using Fiji software (National Institutes of Health, USA). The following primary and secondary antibodies were employed: mouse anti-human Ezrin (Abcam, ab4069, 10 µg/ml); rabbit anti-human ZO-1 (Thermo Fisher Scientific, 61-7300, 2.5 µg/ml); Alexa Fluor 488 anti-mouse IgG (H+L) (Invitrogen, A11001, 8 µg/ml); Alexa Fluor 594 anti-rabbit IgG (H+L) (Invitrogen, A11012, 4 µg/ml). Finally, Phalloidin-iFluor 647 dye (Abcam, ab176759, 1:1000) was used to visualize F-actin.

### Analysis of monolayer thickness, area, and angle to origin

All measurements were performed by a custom script in Fiji ^[31]^. To quantify the monolayer thickness, the confocal stacks obtained from the F-Actin channel were resliced, and at 10% intervals of the total frame width (with a 5% offset from the frame edge) Intensity line plots were drawn vertically to determine the apical and basal edges of the monolayer. For each line plot, the mean grey value was determined and used to apply a threshold where the F-actin signal rose above the background noise. The monolayer was defined as the vertical distance difference between the apical and basal threshold-crossing points. For better comparability, each value was normalized by the mean of the respective experiment. Furthermore, for each measured value, the distance from the center of the artificial drusen was calculated. To quantify the average cell area, the confocal stacks obtained in the F-Actin channel were flattened by Z Projection. Cellpose software^[32]^ was then used to determine the cell outlines and measure cell area, shape factor, and distance to the center of the artificial drusen. For better comparability, each value was normalized by the mean of the respective experiment. To quantify the angle to origin, the absolute angle from the segmented cells in Cellpose was determined. To calculate the “angle to origin”, first we measured the distance and drew a vector towards the center of the artificial drusen. Then, we calculated the scalar product using the vectorized absolute angle.

### Traction force microscopy

Traction force microscopy was performed as described previously^[22,33,34]^. Artificial drusen were induced on a week-old monolayer cultured on a PDH substrate containing fluorescent beads (0.1 µL of 50% solution, Invitrogen FluoSpheres™ 0.1 µm orange, 540/560). Samples were imaged with an Axio Observer 7 microscope (Carl Zeiss) at 20X magnification in brightfield and red fluorescence. 24 h after UV illumination a Z-stack (40 slices, 4 µm step) was taken, and the cells were detached with 3 mL 10% SDS. This was followed by 30 min equilibration time for the hydrogel and a reference Z-stack was recorded. These Z-stacks were projected and aligned using the Template Matching plugin in Fiji. *Top*, *Edge*, and *Outside* regions were analyzed with the Particle Image Velocimetry (PIV) plugin to determine bead displacement, and traction forces were calculated with the Fourier Transform Traction Cytometry (FTTC) plugin to obtain mean and median normal stress magnitudes.

### Statistical Analysis

All statistical analyses and data visualisations were conducted using Prism 10 (GraphPad Software) or Origin 2025 (OriginLab Corporation). Before performing statistical analyses, some data were pre-processed to ensure consistency across experiments and to account for potential variation between replicates. The swelling height was expressed as a percentage and normalised through standardisation to the interval [0, 1] using the built-in normalisation function in Origin 2025. Similarly, the monolayer height values were normalised to the average monolayer height of each respective technical replicate using the same software. During this pre-processing step, the data were also evaluated for potential outliers, which were excluded where justified based on deviation from the experimental range or technical artefacts. The statistical tests performed, the number of independent repetitions (N), and the p-values for each experiment are indicated in the corresponding figure caption. The number of technical replicates (n) per independent replicate of each experiment is shown in the supplementary table 1.

### Declaration of AI use

The preparation of this manuscript has been aided by AI-assisted technologies to improve its readability and language.

## Acknowledgment

We would like to acknowledge all the members of the REMeD group in the past years for their contributions to the discussion of the work. Moreover, we appreciate Adam Breitscheidel’s technical support in graphic design, Gloria’s excellent help in drusen induction, and Sebastian Pape for the help with the Fuji script.

## Author contributions

FR: methodology, software, analysis, investigation, writing, and visualization. CB: methodology. IS: methodology, investigation. TP: investigation. VTK: investigation. AK: investigation. SY: investigation. LDL: supervision, resources. JDR: supervision, resources, funding acquisition, writing, conceptualization, and visualization.

## Data Availability Statement

The data that support the findings of this study are available from the corresponding author upon reasonable request

## Supporting Information

**Supplementary Figure 1:**
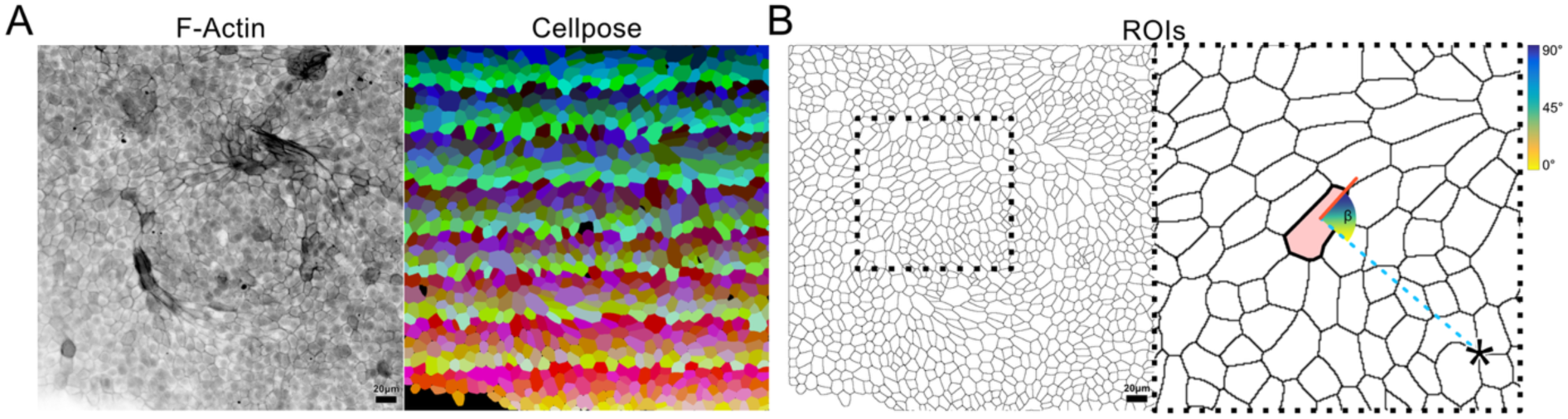
Methodology for quantifying cellular angles to the origin. **A)** Representative maximum intensity projection of F-actin-stained RPE with the Cellpose segmentation output picture. **B)** Representative maximum projection of a ZO-1-stained RPE with the zoomed-in region highlighted by a dashed line. The center of the artificial drusen is marked by an asterisk. The absolute angle of the cell is indicated by the red solid line, while the vector from the center of the highlighted cell to the center of the artificial drusen is indicated by the dotted line. The scalar product between the two vectors is indicated by angle β and lies between 0° and 90°, as shown by a color gradient from yellow to blue.

**Supplementary Figure 2:**
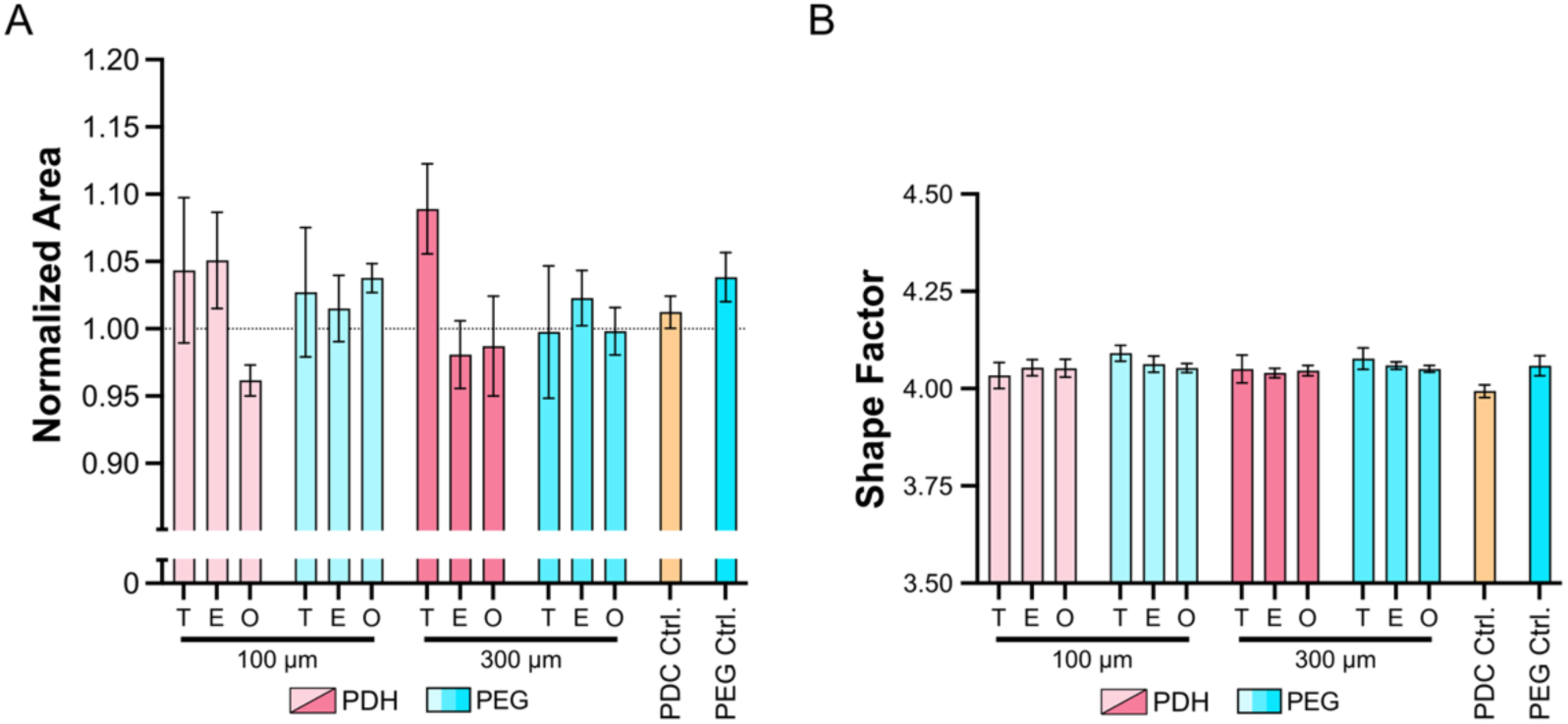
RPE cell areas and shape factors. **A)** Bar plot showing normalized average cell area ± SEM in all conditions. **C)** Bar plot showing the normalized shape factor ± SEM in all conditions.

**Supplementary Figure 3:**
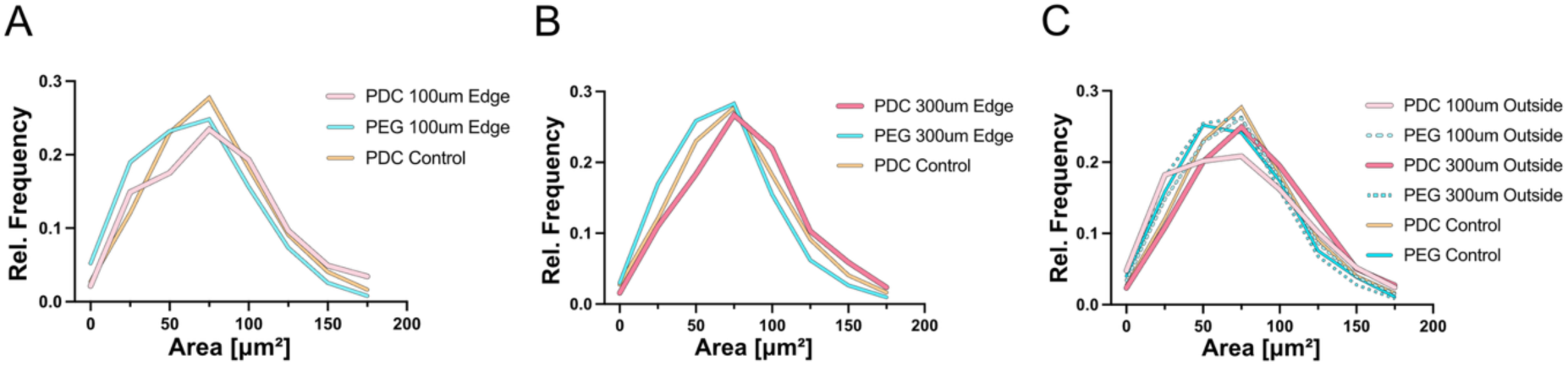
Cell area distribution. Relative frequency distributions of the cell areas for RPE on 100 µm Edge (A) and controls, 300 µm Edge and controls (B), and 100 µm, 300 µm Outside and controls (C).

**Supplementary Table 1.**
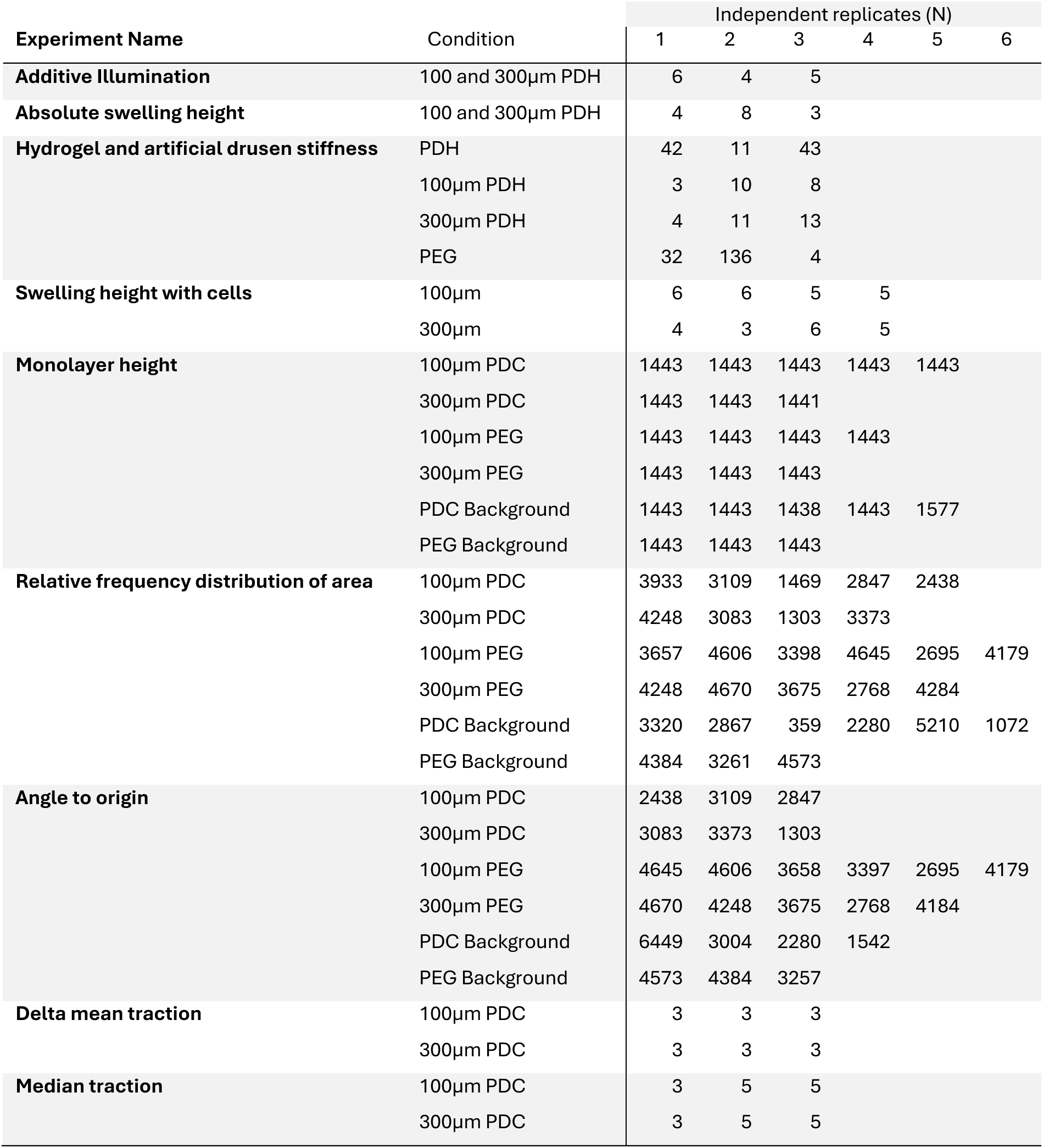
Number of technical replicates per independent replicate.

## References

[1] P. Lu, K. Takai, V. M. Weaver, Z. Werb, Csh Perspect Biol 2011, 3, a005058.

[2] A. N. Kozyrina, T. Piskova, J. Di-Russo, Frontiers Bioeng Biotechnology 2020, 8, 596599.

[3] C. Frantz, K. M. Stewart, V. M. Weaver, J Cell Sci 2010, 123, 4195.

[4] K. B. Jensen, M. H. Little, Stem Cell Rep. 2023, 18, 1255.

[5] B. Artegiani, D. Hendriks, Dev. Cell 2025, 60, 493.

[6] E. L. Fletcher, A. I. Jobling, U. Greferath, S. A. Mills, M. Waugh, T. Ho, R. U. de Iongh, J. A. Phipps, K. A. Vessey, Optom. Vis. Sci. 2014, 91, 878.

[7] S. Chen, N. A. Popp, C.-C. Chan, Expert Rev. Ophthalmol. 2014, 9, 285.

[8] T. J. Heesterbeek, L. Lorés-Motta, C. B. Hoyng, Y. T. E. Lechanteur, A. I. den Hollander, Ophthal Physl Opt 2020, 40, 140.

[9] S. M. Waldstein, W.-D. Vogl, H. Bogunovic, A. Sadeghipour, S. Riedl, U. Schmidt-Erfurth, Jama Ophthalmol 2020, 138, 740.

[10] S. Goverdhan, H. Thomson, A. Lotery, Drug Discov. Today: Dis. Model. 2014, 10, e181.

[11] M. E. Pennesi, M. Neuringer, R. J. Courtney, Mol Aspects Med 2012, 33, 487.

[12] F. Farjood, A. Ahmadpour, S. Ostvar, E. Vargis, J. Biol. Eng. 2020, 14, 13.

[13] K. Yamaguchi, T. Matsuo, F. Shiraga, H. Ohtsuki, Jpn. J. Ophthalmol. 2001, 45, 470.

[14] F. Farjood, E. Vargis, Lab Chip 2018, 18, 3413.

[15] A. M. Kloxin, A. M. Kasko, C. N. Salinas, K. S. Anseth, Science 2009, 324, 59.

[16] A. M. Kloxin, M. W. Tibbitt, K. S. Anseth, Nat. Protoc. 2010, 5, 1867.

[17] A. Buxboim, K. Rajagopal, A. E. X. Brown, D. E. Discher, J. Phys.: Condens. Matter 2010, 22, 194116.

[18] N. R. Richbourg, N. A. Peppas, Prog. Polym. Sci. 2020, 105, 101243.

[19] Y. J. Chuah, Z. T. Heng, J. S. Tan, L. M. Tay, C. S. Lim, Y. Kang, D.-A. Wang, Colloids Surf. B: Biointerfaces 2020, 191, 110995.

[20] A. Wolfel, M. Jin, J. I. Paez, Front. Chem. 2022, 10, 1012443.

[21] R. L. Wilson, G. Swaminathan, K. Ettayebi, C. Bomidi, X.-L. Zeng, S. E. Blutt, M. K. Estes, K. J. Grande-Allen, Tissue Eng. Part C: Methods 2021, 27, 12.

[22] A. N. Kozyrina, T. Piskova, F. Semeraro, I. C. Doolaar, T. Prapty, T. Haraszti, M. Hubert, R. Windoffer, R. E. Leube, A.-S. Smith, J. Di-Russo, EMBO Rep. 2025, 1.

[23] T. Piskova, A. N. Kozyrina, G. Astrauskaitė, M. H. E. Mabrouk, S. Schepl, S. L. S. Yam, R. Ravithas, W. Wagner, M. Vassalli, J. Di-Russo, bioRxiv 2025, DOI 10.1101/2025.05.22.655517.

[24] V. L. Bonilha, S. C. Finnemann, E. Rodriguez-Boulan, J. Cell Biol. 1999, 147, 1533.

[25] V. L. Bonilha, M. E. Rayborn, I. Saotome, A. I. McClatchey, J. G. Hollyfield, Exp. Eye Res. 2006, 82, 720.

[26] D. Bi, J. H. Lopez, J. M. Schwarz, M. L. Manning, Nat Phys 2015, 11, 1074.

[27] A. Nestor-Bergmann, G. Goddard, S. Woolner, O. E. Jensen, Math. Med. Biol.: A J. IMA 2018, 35, 1.

[28] D. T. Tambe, C. C. Hardin, T. E. Angelini, K. Rajendran, C. Y. Park, X. Serra-Picamal, E. H. Zhou, M. H. Zaman, J. P. Butler, D. A. Weitz, J. J. Fredberg, X. Trepat, Nat Mater 2011, 10, 469.

[29] T. P. J. Wyatt, A. R. Harris, M. Lam, Q. Cheng, J. Bellis, A. Dimitracopoulos, A. J. Kabla, G. T. Charras, B. Baum, Proc. Natl. Acad. Sci. 2015, 112, 5726.

[30] Q. Zhang, M. A. Chrenek, S. Bhatia, A. Rashid, S. Ferdous, K. J. Donaldson, H. Skelton, W. Wu, T. R. O. See, Y. Jiang, N. Dalal, J. M. Nickerson, H. E. Grossniklaus, Mol. Vis. 2019, 25, 70.

[31] J. Schindelin, I. Arganda-Carreras, E. Frise, V. Kaynig, M. Longair, T. Pietzsch, S. Preibisch, C. Rueden, S. Saalfeld, B. Schmid, J.-Y. Tinevez, D. J. White, V. Hartenstein, K. Eliceiri, P. Tomancak, A. Cardona, Nat. Methods 2012, 9, 676.

[32] C. Stringer, T. Wang, M. Michaelos, M. Pachitariu, Nat Methods 2021, 18, 100.

[33] M. Vishwakarma, J. D. Russo, D. Probst, U. S. Schwarz, T. Das, J. P. Spatz, Nat Commun 2018, 9, 3469.

[34] J. Di-Russo, J. L. Young, A. Balakrishnan, A. S. Benk, J. P. Spatz, Biomaterials 2019, 192, 171

