## supplementary table 1 for "Photodegradable Hydrogels for On-Demand Modeling of Age-Related Spatiotemporal ECM Deformation"

| Number of technical replicates per independent replicate | | | |  |  |  |  |  |  |
| --- | --- | --- | --- | --- | --- | --- | --- | --- | --- |
|  |  |  | | Independent replicates (N) | | | | | |
|  | Experiment Name | | Condition | 1 | 2 | 3 | 4 | 5 | 6 |
|  | Additive Illumination | 100 and 300µm PDH | | 6 | 4 | 5 |  |  |  |
|  | Absolute swelling height | 100 and 300µm PDH | | 4 | 8 | 3 |  |  |  |
|  | Hydrogel and artificial drusen stiffness | PDH | | 42 | 11 | 43 |  |  |  |
|  |  | 100µm PDH | | 3 | 10 | 8 |  |  |  |
|  |  | 300µm PDH | | 4 | 11 | 13 |  |  |  |
|  |  | PEG | | 32 | 136 | 4 |  |  |  |
|  | Swelling height with cells | 100µm | | 6 | 6 | 5 | 5 |  |  |
|  |  | 300µm | | 4 | 3 | 6 | 5 |  |  |
|  | Monolayer height | 100µm PDC | | 1443 | 1443 | 1443 | 1443 | 1443 |  |
|  |  | 300µm PDC | | 1443 | 1443 | 1441 |  |  |  |
|  |  | 100µm PEG | | 1443 | 1443 | 1443 | 1443 |  |  |
|  |  | 300µm PEG | | 1443 | 1443 | 1443 |  |  |  |
|  |  | PDC Background | | 1443 | 1443 | 1438 | 1443 | 1577 |  |
|  |  | PEG Background | | 1443 | 1443 | 1443 |  |  |  |
|  | Relative frequency distribution of area | 100µm PDC | | 3933 | 3109 | 1469 | 2847 | 2438 |  |
|  |  | 300µm PDC | | 4248 | 3083 | 1303 | 3373 |  |  |
|  |  | 100µm PEG | | 3657 | 4606 | 3398 | 4645 | 2695 | 4179 |
|  |  | 300µm PEG | | 4248 | 4670 | 3675 | 2768 | 4284 |  |
|  |  | PDC Background | | 3320 | 2867 | 359 | 2280 | 5210 | 1072 |
|  |  | PEG Background | | 4384 | 3261 | 4573 |  |  |  |
|  | Angle to origin | 100µm PDC | | 2438 | 3109 | 2847 |  |  |  |
|  |  | 300µm PDC | | 3083 | 3373 | 1303 |  |  |  |
|  |  | 100µm PEG | | 4645 | 4606 | 3658 | 3397 | 2695 | 4179 |
|  |  | 300µm PEG | | 4670 | 4248 | 3675 | 2768 | 4184 |  |
|  |  | PDC Background | | 6449 | 3004 | 2280 | 1542 |  |  |
|  |  | PEG Background | | 4573 | 4384 | 3257 |  |  |  |
|  | Delta mean traction | 100µm PDC | | 3 | 3 | 3 |  |  |  |
|  |  | 300µm PDC | | 3 | 3 | 3 |  |  |  |
|  | Median traction | 100µm PDC | | 3 | 5 | 5 |  |  |  |
|  |  | 300µm PDC | | 3 | 5 | 5 |  |  |  |
