## Supplementary figures and images for "Photodegradable Hydrogels for On-Demand Modeling of Age-Related Spatiotemporal ECM Deformation"

### Supplementary Figure 1

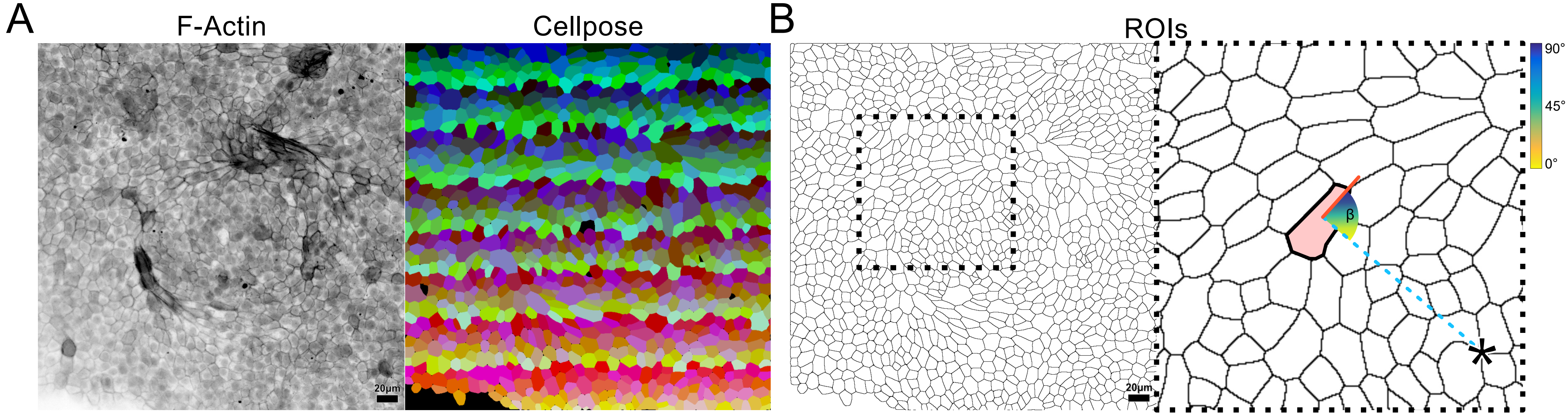

### Supplementary Figure 2

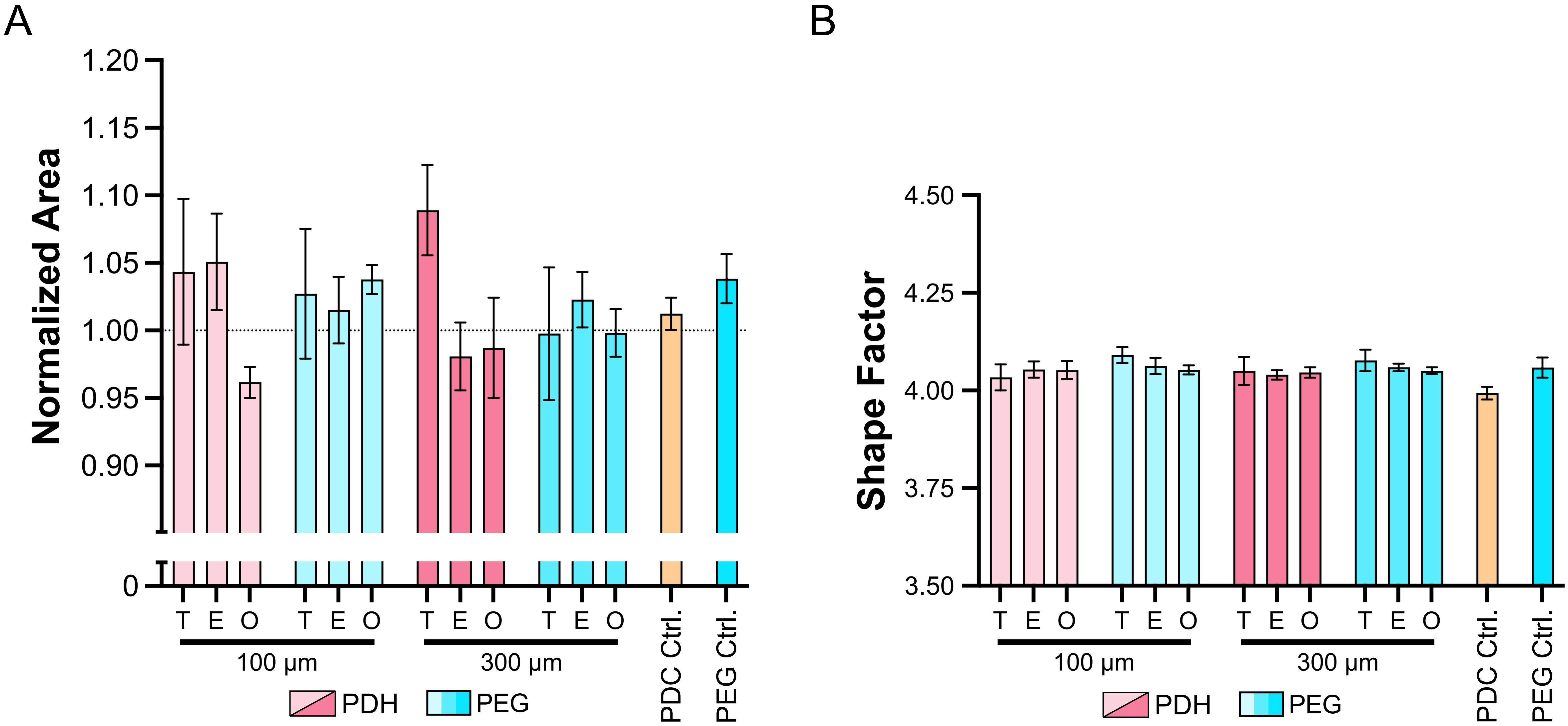

### Supplementary Figure 3

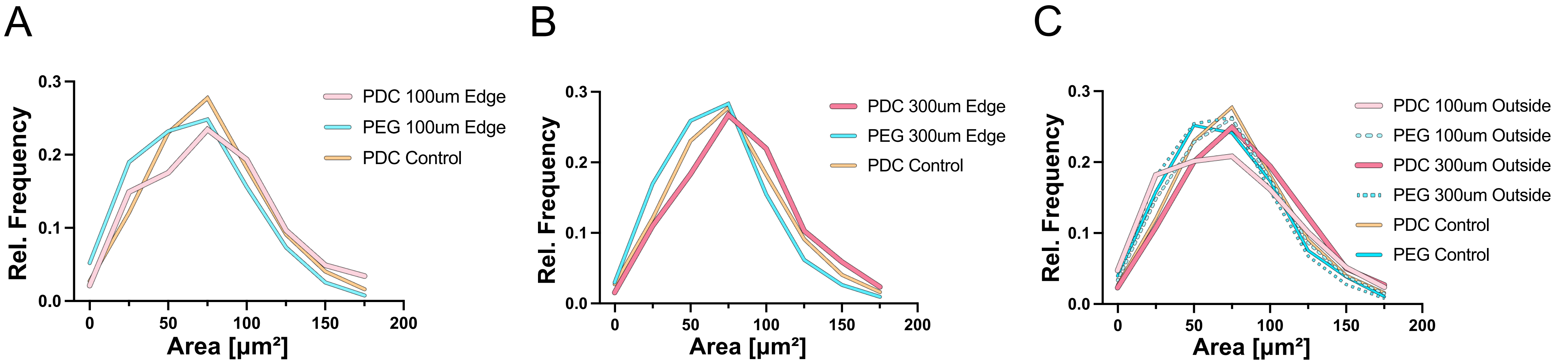
